# Over-Synchrony: Higher Maternal Neuroticism Associates with Stronger Interpersonal Neural Synchrony with Child During Passive and Free Interactions

**DOI:** 10.64898/2026.03.25.714118

**Authors:** Alessandro Carollo, Andrea Bizzego, Dorina Shermadhi, Dagmara Dimitriou, Ilanit Gordon, Gianluca Esposito, Stefanie Hoehl

## Abstract

Interpersonal neural synchrony (INS) in mother–child dyads is often interpreted as a neural marker of relational quality and sensitive caregiving, yet findings on its predictors remain heterogeneous. One possible source of this variability is the diversity of interactional paradigms used in hyperscanning research. This study examined how maternal personality, child temperament, and affective states relate to INS across interaction contexts varying in social interactivity. Thirty-three mother–child dyads (*n* = 20 female children) participated in a functional near-infrared spectroscopy hyperscanning experiment involving passive video co-exposure, a structured cooperative task, and free interaction. Fronto-temporal activity was recorded simultaneously, and INS was computed using wavelet transform coherence. Above-chance levels of INS emerged in inter-brain region combinations primarily involving the mother’s left inferior frontal gyrus (IFG) and the child’s right IFG (adjusted *p*s < 0.030, Cohen’s *d* range = 0.14–0.31). Maternal neuroticism was the only significant predictor of INS, with higher levels associated with increased synchrony during passive video co-exposure (adjusted *p* = 0.012) and free interaction (adjusted *p* = 0.021), but not during the structured game. These findings indicate that maternal dispositional traits shape INS in a context-dependent manner. Notably, the positive association between neuroticism and INS suggests that heightened neural synchrony may reflect over-attunement in more anxious caregivers, rather than optimal coordination. Excessive synchrony may therefore index tightly coupled, over-monitoring interaction dynamics, consistent with models of affiliative vigilance in anxious parenting. Overall, INS may follow a non-linear pattern in which moderate levels are most adaptive, highlighting its flexible, dynamic, and context-sensitive nature.

## 1. Introduction

From the very beginning of life, interactions with caregivers play a fundamental role in shaping human development (Bowlby, 1969). Early bonds with caregivers provide the primary relational context in which children’s socio-cognitive and emotional capacities emerge and develop (Bornstein et al., 2015).

To characterize these early bonds and relational patterns, developmental research has conceptualized the caregiver–child dyad as a functional system, whose dynamics are shaped by the states and actions of both partners (Feldman, 2007). This systemic perspective is traditionally illustrated by the Still-Face Paradigm (Tronick et al., 1978), which shows how brief disruptions in caregiver responsiveness affect infants’ emotional regulation and interactive behavior (Haley and Stansbury, 2003; Lowe et al., 2012).

When adopting a systemic view, one emergent property of the caregiver-child system is a pattern of coordination at multiple biological and behavioral levels (Carollo et al., 2021; Feldman, 2017). This property, known as interpersonal synchrony, refers to the temporal coordination of events between interacting individuals, and it emerges during social interactions across the lifespan (Feldman, 2017). From a physical and biological perspective, synchronization can be understood as a low-cost dynamical regime in which coupled systems require reduced regulatory and computational effort to maintain coordination (e.g., Friston et al., 2006; Friston, 2010; Strogatz, 2000). By aligning their temporal dynamics, interacting individuals may reduce uncertainty and stabilize the dyadic system, thereby supporting efficient interpersonal regulation and information exchange (De Reus et al., 2021). In caregiver-child dyads, interpersonal synchrony tends to associate with a healthy mother, typical development, and more positive child cognitive and behavioral outcomes (Leclère et al., 2014).

The interpersonal synchrony framework initially arose from the study of behavioral coordination (Feldman, 2006; Stern et al., 1977). However, recent advances in neuroscience have made it possible to investigate coordination at the neural level between interacting individuals, thereby adding a further level of information (Carollo and Esposito, 2024; De Felice et al., 2025; Montague et al., 2002). In particular, the simultaneous recording of brain activity from multiple individuals (i.e., hyperscanning) has shown that caregiver–child brain activity synchronizes during social interactions (Hoehl et al., 2025). This phenomenon, termed interpersonal neural synchrony (INS), emerges across different tasks, ranging from passive video co-watching to active social interactions with a shared goal (Azhari et al., 2019; Nguyen et al., 2020).

Hyperscanning studies indicate that INS reflects variations in the behaviors and internal states of the individual members of the caregiver–child system. For example, an electroencephalography (EEG) hyperscanning study by Endevelt-Shapira and Feldman (2023) showed that distinct parenting styles are differentially related to INS during face-to-face interactions between mothers and their 5–12-month-old infants. Specifically, greater maternal sensitivity has been linked to increased mother–infant neural synchrony, whereas higher maternal intrusiveness has been associated with reduced inter-brain coordination. Consistent with these findings, functional near-infrared spectroscopy (fNIRS) studies have shown that INS is associated with interaction quality and collaborative problem-solving during cooperative mother–child tasks, particularly within fronto-temporal regions (Nguyen et al., 2020; Reindl et al., 2018). These fronto-temporal regions include areas such as the inferior frontal gyrus (IFG) and the temporoparietal junction (TPJ), which are consistently implicated in social cognition and interactive coordination. The IFG is associated with action understanding, imitation, and the integration of sensorimotor signals during social interaction (Iacoboni and Dapretto, 2006). Moreover, the IFG appears to encode for the social gap between self and others and to coordinate responses to increase interpersonal closeness (Kanterman and Shamay-Tsoory, 2025; Ravreby et al., 2022; Shamay-Tsoory et al., 2019). The TPJ has been linked to perspective-taking and mentalizing processes that support the interpretation of others’ intentions and behaviors (Saxe, 2010). Additional evidence suggests that greater prefrontal INS between parents and children may play a role in child’s socio-emotional development and emotional regulation (Reindl et al., 2018). Higher levels of parental stress have been associated with reduced INS across interactive contexts (Azhari et al., 2019; Nguyen et al., 2020). Additionally, Morgan et al. (2023) showed that INS is time-linked to positive affective state matching in mother-child dyads. Nevertheless, associations between parental behavior, attachment representations, and INS are not uniformly positive. For instance, Nguyen et al. (2024) reported higher frontal INS in dyads characterized by insecure maternal attachment representations. The authors suggest that this pattern may reflect heightened maternal efforts to coordinate with and anticipate the child’s behavior, supporting the idea of a potential optimal balance between tendencies toward synchrony and segregation in everyday social interactions (Beebe and McCrorie, 2010; Gordon et al., 2024).

However, the relationship between these variables and INS in mother–child dyads has been examined across highly heterogeneous interactive contexts, ranging from passive conditions to more active forms of engagement (Azhari et al., 2019; Nguyen et al., 2020). Such contextual heterogeneity may partly account for inconsistencies in the strength and direction of associations between caregivers’ and children’s dispositional or situational characteristics and levels of INS. Indeed, the interactive context plays a key role in modulating INS (Carollo et al., 2025b), likely reflecting the fact that different tasks place distinct demands on interpersonal coordination and therefore require different degrees of INS. To fill this gap, the current work investigates the relationship between mother-child dispositional and situational traits and INS across different levels of social interactivity. The study focuses on preschool-aged children (5 years old), a developmental period characterized by substantial advances in socio-cognitive abilities. At this age, children are increasingly capable of engaging in structured and reciprocal social interactions while still relying on caregivers as primary co-regulatory partners. The data analyzed in the present study are drawn from a larger fNIRS hyperscanning project investigating how interpersonal closeness and social interactivity levels modulate INS (Carollo et al., 2025b). In the study, we expected that individual dispositional and situational factors would be associated with the degree of INS observed during mother–child interaction. In particular, maternal personality traits, children’s temperament, and their affect during the experiment were hypothesized to be reflected in the extent to which dyads synchronized at the neural level during social engagement. Moreover, given that different interactive contexts impose distinct demands on interpersonal coordination, we expected these associations to vary as a function of the level of social interactivity. However, given the heterogeneity of the existing literature, no specific directional hypotheses were formulated regarding these associations.

## 2. Methods

### 2.1. Study design

The present study is part of a larger project investigating how interpersonal closeness and levels of social interactivity modulate INS in a sample of 142 adult–adult and adult–child dyads (Carollo et al., 2025b). To examine the role of interpersonal closeness, the study included three types of dyads: close friends, romantic partners, and mother–child dyads. The current manuscript focuses exclusively on the mother–child dyads data, which were collected at the University of Vienna, in Austria. In addition to the interaction tasks, mothers and children completed a series of pre- and post-interaction questionnaires assessing dispositional (i.e., mother’s personality and child’s temperament) and situational (i.e., affect) characteristics.

All dyads participated in three interaction contexts that differed in levels of social interactivity: passive video co-exposure, a structured rule-based cooperative task, and an unstructured verbal interaction. These activities were chosen to capture a *continuum* of interactivity, ranging from more controlled, low-engagement situations to spontaneous and highly interactive exchanges. Throughout the experiment, brain activity from both members of each dyad was recorded simultaneously using an fNIRS hyperscanning approach.

The project received ethical approval from the Ethics Committees of the University of Trento (ref. 2022–052) and the University of Vienna (ref. 00732), where the data were collected. All procedures were conducted in accordance with the Declaration of Helsinki. Written informed consent was obtained from all adult participants, while consent for child participants was provided by their primary caregivers.

### 2.2. Participants

A total of 33 mother–child dyads participated in the study, including 20 female children. Mothers had a mean age of 38.59 years (*SD* = 4.66), and children had a mean age of 5 years, 6 months, and 18 days (*SD* = 3 months, 22 days). Two dyads were excluded due to technical issues during data collection. The demographic characteristics of the sample are reported in Table 1.

**Table 1:** Demographic information and descriptive statistics of the sample.

| Demographic variable | Value |
| --- | --- |
| Gestational age [weeks] | 40.08 |
| Language household: monolingual [frequency] | 18 |
| Language household: multilingual [frequency] | 13 |
| Language household: missing [frequency] | 2 |
| Socioeconomic status [1-10] | 6.77 |

Inclusion criteria specified that participants were mothers of 5-year-old children. All participants reported no history of medical or neurological disorders, particularly conditions that might influence blood oxygenation, as well as developmental conditions.

Mother–child dyads were recruited via an existing volunteer registry at the University of Vienna. Each dyad received €6 in compensation (equivalent to two round-trip public transport tickets in Vienna), and each child was given a small toy (i.e., a LEGO set worth €6).

### 2.3. Questionnaires

Prior to the experimental procedure, mothers completed the Big Five Inventory–10 (BFI-10) and the Children’s Behavior Questionnaire–Very Short Form (CBQ-VSF). The BFI-10 was used to assess maternal personality across the five major dimensions, while the CBQ-VSF measured child temperament and well-being. Mothers also completed the Positive and Negative Affect Schedule (PANAS) to assess their own and their child’s affective state both before and after the experimental session.

#### 2.3.1. Big Five Inventory

To assess the mother’s personality, the German version of the BFI-10 was used (Rammstedt and John, 2007). The BFI-10 assesses the five major personality dimensions (i.e., Agreeableness, Conscientiousness, Extraversion, Neuroticism, and Openness to Experience) using two items per trait. Responses were provided on a 5-point Likert scale (1 = strongly disagree, 5 = strongly agree). Composite scores were calculated as the mean of the relevant items. Cronbach’s alpha values were low for some traits, particularly Agreeableness (*α* = 0.08) and Conscientiousness (*α* = 0.42), likely reflecting the limited number of items per dimension. In contrast, Extraversion (*α* = 0.80), Neuroticism (*α* = 0.78), and Openness to Experience (*α* = 0.71) showed acceptable internal consistency. As commonly reported for ultra-short personality measures, lower reliability estimates are expected due to the restricted item pool and were retained for the analysis for completeness.

#### 2.3.2. Children’s Behavior Questionnaire

Child temperament was assessed using the German version of the CBQ-VSF (Putnam and Rothbart, 2006). The questionnaire includes 36 items covering three broad dimensions: Effortful Control, Negative Affectivity, and Surgency/Extraversion. Mothers rated how accurately each item described their child’s typical reactions across situations during the previous six months on a 7-point Likert scale (1 = extremely untrue, 7 = extremely true). Scale scores were computed as the mean of the items corresponding to each dimension. Cronbach’s alpha values were acceptable for Effortful Control (*α* = 0.82), Negative Affectivity (*α* = 0.78), and Surgency/Extraversion (*α* = 0.79).

#### 2.3.3. Positive and Negative Affect Schedule

Maternal and child affect were assessed before and after the experiment using the German version of the PANAS (Breyer and Bluemke, 2016; Watson et al., 1988). The PANAS is a 20-item self-report measure comprising two dimensions: Positive Affect, reflecting pleasurable engagement and high activation, and Negative Affect, reflecting general distress and unpleasurable engagement. Mothers rated the extent to which they and their child experienced each affective state on a 5-point Likert scale (1 = very slightly or not at all, 5 = extremely) before and after the experimental session. Positive Affect and Negative Affect scores were computed by summing the relevant items, with higher scores indicating greater levels of positive and negative affect, respectively. Internal consistency was acceptable for maternal Positive Affect (pre: *α* = 0.88; post: *α* = 0.90) and Negative Affect (pre: *α* = 0.95; post: *α* = 0.78). Similarly, child Positive Affect (pre: *α* = 0.89; post: *α* = 0.91) and Negative Affect (pre: *α* = 0.95; post: *α* = 0.82) demonstrated acceptable reliability. Additionally, change scores were computed for both Positive and Negative Affect by subtracting pre-experiment scores from post-experiment scores.

### 2.4. Experimental tasks

Baseline neural activity was assessed through a one-minute resting-state recording at the beginning of the experimental session, which was repeated between tasks. Members of each dyad were seated on separate chairs approximately 60–80 cm apart, and this spatial arrangement was maintained throughout all experimental conditions.

The experimental procedure comprised three tasks: passive video co-exposure, a structured rule-based cooperative game, and an unstructured verbal interaction. These tasks were designed to examine how varying levels of social interactivity influence INS, spanning from highly structured to more spontaneous and less constrained forms of engagement.

In the video co-exposure condition, dyads watched a 3-minute, language-free animated video presented on a monitor in front of them. They were instructed to focus on the video and refrain from interacting with one another. This condition allowed the assessment of INS driven by a shared sensory experience within a social setting, while minimizing the influence of direct interpersonal interaction.

In the rule-based cooperative condition, dyads engaged in a 5-minute game of cooperative Jenga. A Jenga tower was placed on a table between the participants, who were instructed to follow standardized rules: taking turns, removing one block at a time, and placing it on top of the tower. To maintain comparable task difficulty across dyads, participants were asked to remove only one block per layer, starting from the bottom. If the tower collapsed, they were instructed to rebuild it and continue playing, ensuring a consistent task duration across dyads.

In the free verbal interaction condition, participants sat facing one another and engaged in spontaneous conversation for 5 minutes. This unstructured task was intended to elicit flexible and less predictable exchanges characteristic of naturalistic, reciprocal communication.

The order of the experimental conditions was randomized across dyads to control for potential order effects. Resting-state data were not included in the analyses reported in the present study.

### 2.5. Acquisition of neural data

Brain activity was recorded throughout all experimental tasks using an fNIRS hyperscanning setup with a NIRSport system (NIRx Medical Technologies LLC) at a sampling rate of 7.81 Hz. Each participant wore a cap equipped with eight LED sources (wavelengths: 760 and 850 nm) and eight detectors, arranged to measure hemodynamic activity over bilateral inferior frontal gyrus (IFG) and temporoparietal junction (TPJ), consistent with previous research (e.g., Nguyen et al., 2020, 2021). A schematic representation of the optode layout is provided in Figure 1.

**Figure 1:**
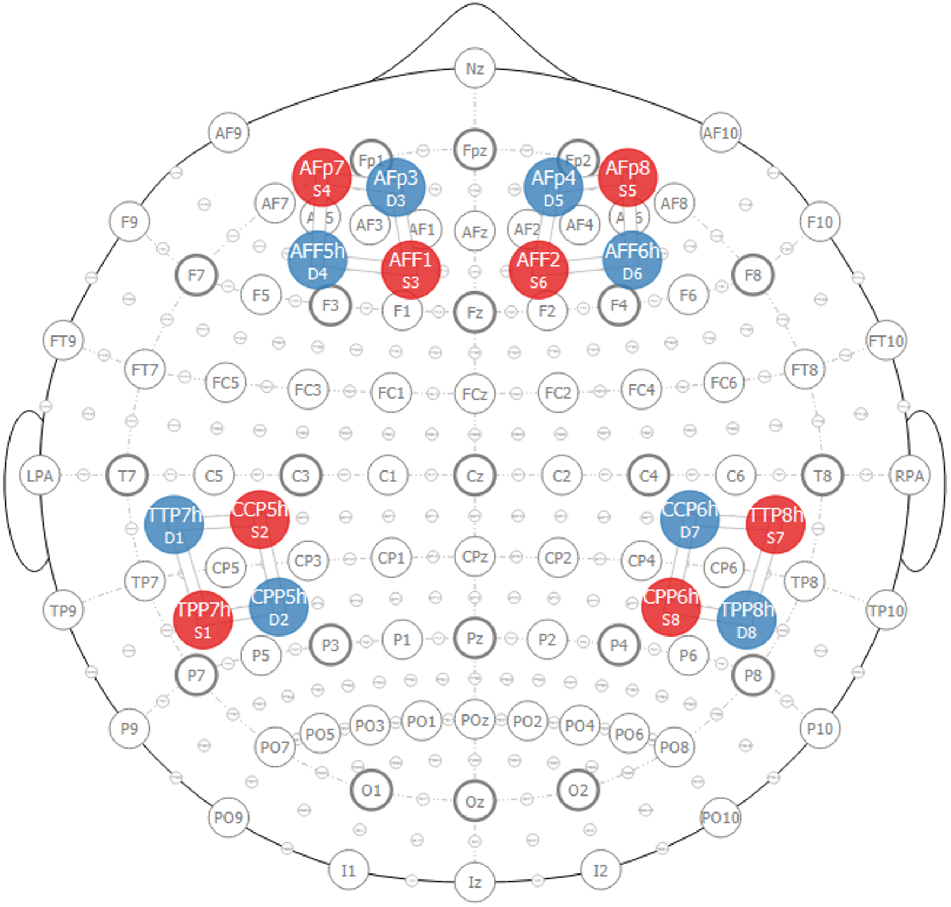
Optode configuration used for functional near-infrared spectroscopy (fNIRS) hyperscanning. Each participant wore a cap with eight light sources and eight detectors, forming 16 measurement channels. Optodes were arranged to record hemodynamic activity over bilateral inferior frontal gyrus and temporoparietal junction.

The cap configuration comprised 16 fNIRS channels per participant, with a source–detector separation of approximately 3 cm. Channels were grouped into four regions of interest (ROIs): left and right IFG and left and right TPJ. The optode arrangement followed configurations used in previous adult–child hyperscanning studies conducted in the same lab (Nguyen et al., 2020, 2021). Channel placement was guided by the international 10–20 EEG system.

### 2.6. Processing of neural data

Pre-processing and synchrony analyses were conducted using *pyphysio* (Bizzego et al., 2019), following the same pipeline adopted in the main project (Carollo et al., 2025b). Raw fNIRS intensity signals were first converted into changes in optical density. To ensure comparable data length across experimental conditions, only the first 3 minutes of each task were included in the analyses. Signal quality was evaluated using a convolutional neural network trained to classify fNIRS segment quality (Bizzego et al., 2021, 2022). Data were then resampled at 10 Hz and converted into concentrations of oxygenated (HbO) and deoxygenated hemoglobin (HbR) using the modified Beer–Lambert Law (Delpy et al., 1988). Consistent with previous work (e.g., Lim et al., 2024a,b), the main statistical analyses focused on HbO signals (see Supplementary Materials for analyses based on HbR).

Temporal autocorrelations and physiological noise were addressed by whitening the signals using an autoregressive model (Barker et al., 2013). This approach has been shown to reduce false discovery rates when estimating coherence between time series (Santosa et al., 2017).

### 2.7. Interpersonal neural synchrony

The current study followed the procedure used in Carollo et al. (2025b), INS was computed on a channel-by-channel basis between dyad members using wavelet transform coherence (WTC) (Grinsted et al., 2004). WTC estimates the coherence between time series across both temporal and frequency domains and captures relationships that occur with zero phase lag as well as delayed (phase-shifted) coupling (Nguyen et al., 2020). This approach enables a detailed characterization of dynamic coherence patterns in brain activity.

For each pair of good-quality channels in each experimental condition, WTC was computed across a frequency range of 0.01–0.20 Hz, with steps of 0.01 Hz, in line with previous studies (Carollo et al., 2025c; Lim et al., 2024a,b). Final WTC values were obtained by averaging coherence estimates across the full frequency range. This strategy avoided privileging specific frequency bands in the absence of *a priori* hypotheses regarding frequencies of interest (Carollo et al., 2025a,b,c).

To estimate a level of INS independent of direct interaction, WTC was also computed for non-interacting surrogate dyads. These surrogate dyads were created by randomly pairing participants from different original dyads, while preserving the adult-child composition of the dyads. Following Carollo et al. (2025b,c), a single permutation of dyads was performed, yielding the same number of observations as in the true dyads.

To ensure consistency in task context and channel correspondence, surrogate pairs were constructed by matching data segments from the same channel and experimental condition across participants. For instance, fNIRS data during the video co-exposure task for participant A (dyad 001) were paired with data from participant B (dyad 002) recorded in the same condition. Synchrony observed in surrogate dyads was assumed to primarily reflect stimulus-driven coupling, thereby providing a baseline against which synchrony in true dyads could be compared to obtain co-presence effects on INS (Golland et al., 2015).

### 2.8. Data analysis

For the current study, we divided the analysis plan into three main parts, with all analyses conducted using linear mixed-effects models (Bates et al., 2005).

For each combination of ROIs across the two participants (e.g., mother left IFG – child right TPJ), we fitted a linear mixed-effects model with WTC as the dependent variable and dyad type (true vs. surrogate) as the fixed effect. Random effects were included to account for multiple sources of non-independence. The experimental condition was modeled to capture condition-specific variance, and dyad ID accounted for repeated measurements within dyads. In addition, variability associated with the pairing of channels across brains was modeled at two complementary levels (as in Carollo et al., 2025b). A channel-pair ID defined independently of participant order treated each pair of ROIs as the same entity, regardless of which member contributed each channel (e.g., mother channel 1 – child channel 2 and child channel 1 – mother channel 2 shared the same identifier), capturing variability related to the intrinsic tendency of specific ROI pairs to exhibit INS. A second channel-pair ID defined as order-dependent treated each ordered configuration as distinct (e.g., mother channel 1 – child channel 2 and child channel 1 – mother channel 2 had different identifier), capturing variability linked to directional configurations across brains and potential role- or brain-specific asymmetries. To control for multiple comparisons across ROI combinations, significance levels were adjusted using a Bonferroni correction (*α* = 0.05/16 = 0.003).

ROI combinations that were sensitive to co-presence and showed a significant hyperscanning effect were then carried forward to subsequent analyses examining associations between INS and dispositional and situational traits at two complementary levels.

At the first level, we performed global analyses in which data from all selected ROI combinations were modeled together, with ROI included as a random effect to account for variability across inter-brain connections. WTC scores served as the dependent variable, while subscale scores from the BFI and CBQ, as well as post-pre changes in PANAS scores, were entered as predictors. The models further included experimental condition, dyad ID, and both order-independent and order-dependent channel-pair IDs as random effects. This approach allowed us to estimate associations that generalize across the inter-brain network while accounting for connection-specific variability.

At the second level, we examined whether these associations varied as a function of the experimental condition. To this end, we fitted three separate linear mixed-effects models, one for each condition (Bonferroni-adjusted *α* = 0.05/3 = 0.017). In each model, WTC scores were entered as the dependent variable, dispositional and situational traits as fixed effects, and dyad ID, along with order-independent and order-dependent channel-pair IDs, as random effects.

## 3. Results

### 3.1. Hyperscanning effect: true versus surrogate dyads

In the first part of the analysis, we identified the combinations of ROIs between mother and child that were sensitive to co-presence. To this end, we fitted a linear mixed-effects model for each ROI combination, with WTC scores as the dependent variable and dyad type as the predictor. A statistically significant hyperscanning effect emerged for the following ROI pairs: mother left IFG – child left IFG, mother left IFG – child right IFG, mother left IFG – child right TPJ, mother right IFG – child right IFG, and mother left TPJ – child right IFG (all Bonferroni-adjusted *p*s *<* 0.031, Cohen’s *d* range = 0.14–0.31). All results are summarized in Table 2. These findings suggest that co-presence selectively enhances INS within a frontotemporal network primarily involving the mother’s left IFG and the child’s right IFG.

**Table 2:** Linear mixed-effects model results comparing true and surrogate mother–child dyads across inter-brain region pairs. Estimates reflect differences in interpersonal neural synchrony between true and randomly paired surrogate dyads for each mother–child region of interest combination (IFG = inferior frontal gyrus; TPJ = temporo-parietal junction). Positive estimates indicate greater synchrony in true dyads relative to surrogate pairs. Reported statistics include fixed-effect estimates, standard errors (SE), degrees of freedom (df), *t*-values, unadjusted *p*-values, multiple-comparison–adjusted *p*-values, and effect sizes where applicable. Significant effects after correction are highlighted in bold. (** Adjusted *p*-value *<* 0.010; *** Adjusted *p*-value *<* 0.001)

| Fixed Effect | Estimate | SE | df | t-value | p-value | Adjusted p-value | Cohen's *d* 95% CI |
| --- | --- | --- | --- | --- | --- | --- | --- |
| **Mother left IFG - Child left IFG** |  |  |  |  |  |  |  |
| True dyads | 0.0031 | 0.001 | 2211 | 5.94 | < 0.001 | < **0.001** *** | 0.25 [0.17, 0.34] |
| **Mother left IFG - Child right IFG** |  |  |  |  |  |  |  |
| True dyads | 0.0036 | 0.001 | 2221 | 7.19 | < 0.001 | < **0.001** *** | 0.31 [0.22, 0.39] |
| **Mother left IFG - Child left TPJ** |  |  |  |  |  |  |  |
| True dyads | 0.0018 | 0.001 | 2142 | 3.30 | 0.001 | **0.031** * | 0.14 [0.06, 0.23] |
| **Mother left IFG - Child right TPJ** |  |  |  |  |  |  |  |
| True dyads | 0.0030 | 0.001 | 2227 | 5.32 | < 0.001 | < **0.001** *** | 0.23 [0.14, 0.31] |
| **Mother right IFG - Child left IFG** |  |  |  |  |  |  |  |
| True dyads | 0.0011 | 0.001 | 2045 | 1.89 | > 0.050 | > 0.050 | / |
| **Mother right IFG - Child right IFG** |  |  |  |  |  |  |  |
| True dyads | 0.0026 | 0.001 | 2051 | 4.90 | < 0.001 | < **0.001** *** | 0.22 [0.13, 0.30] |
| **Mother right IFG - Child left TPJ** |  |  |  |  |  |  |  |
| True dyads | 0.0013 | 0.001 | 2014 | 2.23 | 0.026 | > 0.050 | / |
| **Mother right IFG - Child right TPJ** |  |  |  |  |  |  |  |
| True dyads | 0.0017 | 0.001 | 2053 | 2.97 | 0.003 | > 0.050 | / |
| **Mother left TPJ - Child left IFG** |  |  |  |  |  |  |  |
| True dyads | -0.0002 | 0.001 | 2191 | -0.37 | > 0.050 | > 0.050 | / |
| **Mother left TPJ - Child right IFG** |  |  |  |  |  |  |  |
| True dyads | 0.0019 | 0.001 | 2209 | 3.67 | < 0.001 | **0.008** ** | 0.16 [0.07, 0.24] |
| **Mother left TPJ - Child left TPJ** |  |  |  |  |  |  |  |
| True dyads | -0.0001 | 0.001 | 2147 | -0.12 | > 0.050 | > 0.050 | / |
| **Mother left TPJ - Child right TPJ** |  |  |  |  |  |  |  |
| True dyads | -0.0018 | 0.001 | 2201 | -3.12 | 0.0022 | > 0.050 | / |
| **Mother right TPJ - Child left IFG** |  |  |  |  |  |  |  |
| True dyads | 0.0015 | 0.001 | 1942 | 2.82 | 0.005 | > 0.050 | / |
| **Mother right TPJ - Child right IFG** |  |  |  |  |  |  |  |
| True dyads | 0.0018 | 0.001 | 1921 | 3.13 | 0.002 | > 0.050 | / |
| **Mother right TPJ - Child left TPJ** |  |  |  |  |  |  |  |
| True dyads | 0.0011 | 0.001 | 1881 | 1.79 | > 0.050 | > 0.050 | / |
| **Mother right TPJ - Child right TPJ** |  |  |  |  |  |  |  |
| True dyads | 0.0007 | 0.001 | 1935 | 1.09 | > 0.050 | > 0.050 | / |

### 3.2. Association between dispositional and situational traits and interpersonal neural synchrony

After identifying the ROI combinations sensitive to co-presence, we investigated how dispositional and situational traits were associated with levels of INS in these connections. To this end, we fitted a linear mixed-effects model across ROI combinations, with WTC scores as the dependent variable and subscale scores indexing maternal personality traits, child temperament, and changes in affect as predictors. The model included experimental condition, ROI combination, order-dependent and order-independent channel-pair identifiers, and dyad ID as random effects.

Of all predictors entered in the model, only maternal neuroticism was significantly associated with WTC levels, with higher scores predicting increased INS (*β* = 0.003, *p* = 0.032). The other predictors did not show a statistically significant association with INS (*p*s *>* 0.050). The model explained a small but meaningful portion of the variance, with a marginal *R*^2^ of 0.04. The full results of this model are reported in Table 3.

**Table 3:**
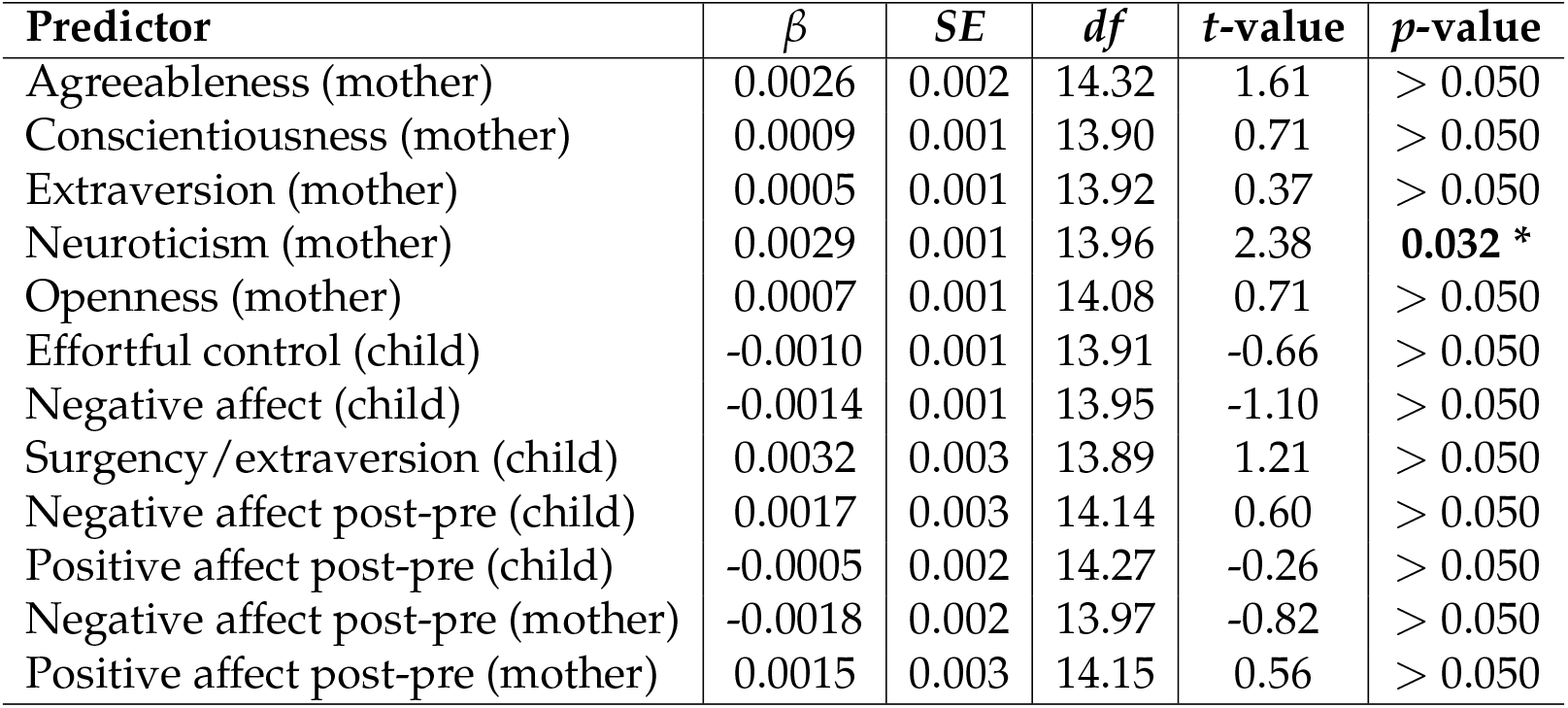
Linear mixed-effects model examining dispositional and situational predictors of interpersonal neural synchrony. Maternal personality traits, child temperament dimensions, and post-pre changes in affect for both mother and child were entered as predictors of global interpersonal neural synchrony computed across region of interest pairs that previously demonstrated a significant hyperscanning effect. Reported statistics include regression coefficients (*β*), standard errors (SE), degrees of freedom (df), *t*-values, and associated *p*-values. Significant predictors are highlighted in bold. (* *p*-value *<* 0.050)

| Predictor | $\beta$ | SE | df | t-value | p-value |
| --- | --- | --- | --- | --- | --- |
| Agreeableness (mother) | 0.0026 | 0.002 | 14.32 | 1.61 | > 0.050 |
| Conscientiousness (mother) | 0.0009 | 0.001 | 13.90 | 0.71 | > 0.050 |
| Extraversion (mother) | 0.0005 | 0.001 | 13.92 | 0.37 | > 0.050 |
| Neuroticism (mother) | 0.0029 | 0.001 | 13.96 | 2.38 | **0.032 \*** |
| Openness (mother) | 0.0007 | 0.001 | 14.08 | 0.71 | > 0.050 |
| Effortful control (child) | -0.0010 | 0.001 | 13.91 | -0.66 | > 0.050 |
| Negative affect (child) | -0.0014 | 0.001 | 13.95 | -1.10 | > 0.050 |
| Surgency/extraversion (child) | 0.0032 | 0.003 | 13.89 | 1.21 | > 0.050 |
| Negative affect post-pre (child) | 0.0017 | 0.003 | 14.14 | 0.60 | > 0.050 |
| Positive affect post-pre (child) | -0.0005 | 0.002 | 14.27 | -0.26 | > 0.050 |
| Negative affect post-pre (mother) | -0.0018 | 0.002 | 13.97 | -0.82 | > 0.050 |
| Positive affect post-pre (mother) | 0.0015 | 0.003 | 14.15 | 0.56 | > 0.050 |

### 3.3. Association between dispositional and situational traits and interpersonal neural synchrony across social interactivity levels

When examining these effects within individual experimental conditions, maternal neuroticism emerged as the only statistically significant predictor of INS in both the video co-exposure condition (*β* = 0.006, adjusted *p* = 0.011) and the free interaction condition (*β* = 0.004, adjusted *p* = 0.020). The model predicting INS during video co-exposure showed a marginal *R*^2^ of 0.12, whereas the model for the free interaction condition showed a marginal *R*^2^ of 0.07. None of the other predictors were significantly associated with INS in any condition (all adjusted *p*s *>* 0.050). Results for each experimental condition are reported in Table 4.

**Table 4:**
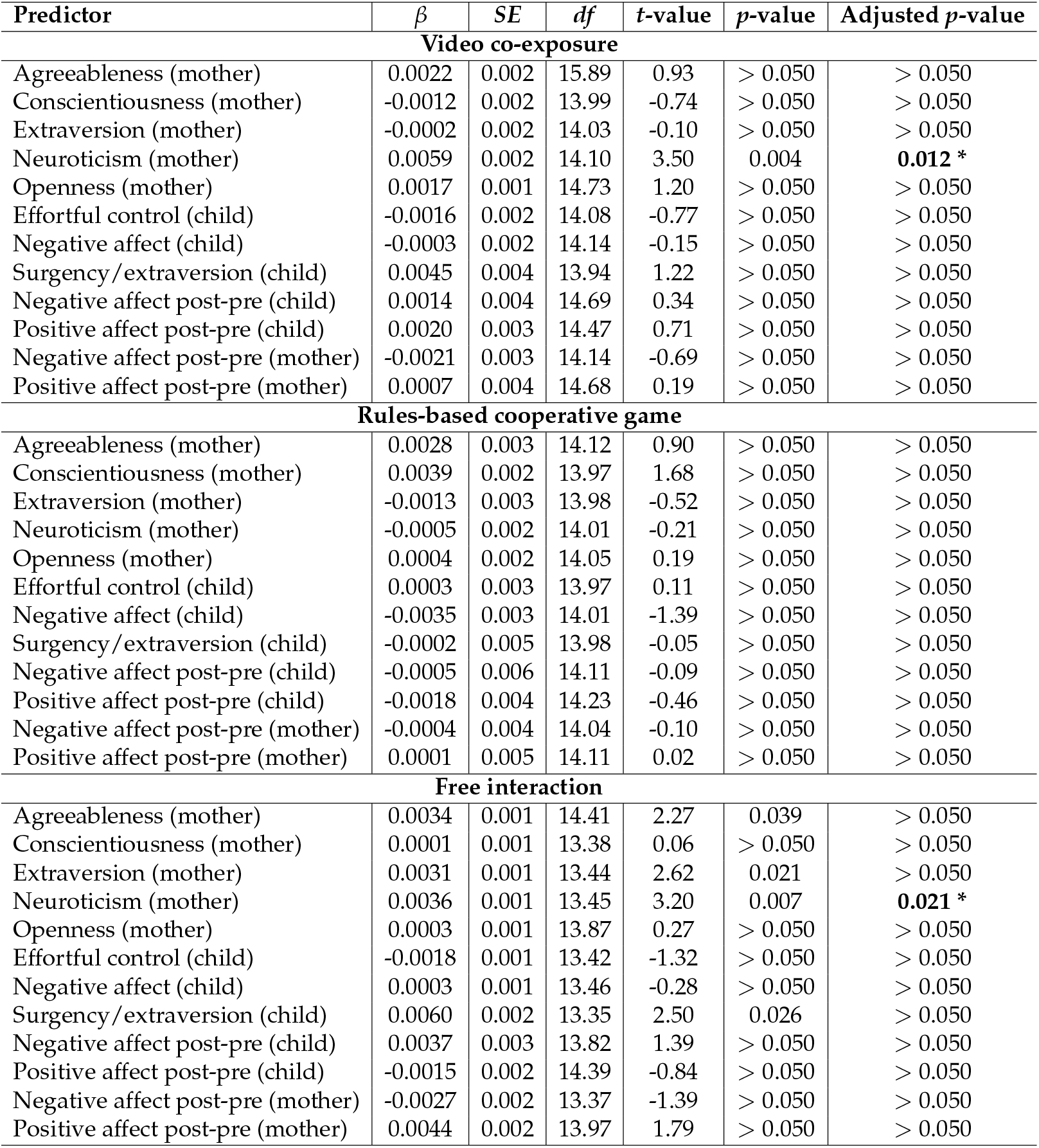
Linear mixed-effects model results examining whether maternal personality traits, child temperament dimensions, and post-pre affective changes predict interpersonal neural synchrony across region of interest pairs previously identified as showing a hyperscanning effect, separately for each experimental condition. Reported statistics include regression coefficients (*β*), standard errors (SE), degrees of freedom (df), *t*-values, unadjusted *p*-values, and multiple-comparison–adjusted *p*-values. Significant predictors after correction are highlighted in bold. (* Adjusted *p*-value *<* 0.050)

| Predictor | $\beta$ | SE | df | t-value | p-value | Adjusted p-value |
| --- | --- | --- | --- | --- | --- | --- |
| **Video co-exposure** |  |  |  |  |  |  |
| Agreeableness (mother) | 0.0022 | 0.002 | 15.89 | 0.93 | > 0.050 | > 0.050 |
| Conscientiousness (mother) | -0.0012 | 0.002 | 13.99 | -0.74 | > 0.050 | > 0.050 |
| Extraversion (mother) | -0.0002 | 0.002 | 14.03 | -0.10 | > 0.050 | > 0.050 |
| Neuroticism (mother) | 0.0059 | 0.002 | 14.10 | 3.50 | 0.004 | **0.012 \*** |
| Openness (mother) | 0.0017 | 0.001 | 14.73 | 1.20 | > 0.050 | > 0.050 |
| Effortful control (child) | -0.0016 | 0.002 | 14.08 | -0.77 | > 0.050 | > 0.050 |
| Negative affect (child) | -0.0003 | 0.002 | 14.14 | -0.15 | > 0.050 | > 0.050 |
| Surgency/extraversion (child) | 0.0045 | 0.004 | 13.94 | 1.22 | > 0.050 | > 0.050 |
| Negative affect post-pre (child) | 0.0014 | 0.004 | 14.69 | 0.34 | > 0.050 | > 0.050 |
| Positive affect post-pre (child) | 0.0020 | 0.003 | 14.47 | 0.71 | > 0.050 | > 0.050 |
| Negative affect post-pre (mother) | -0.0021 | 0.003 | 14.14 | -0.69 | > 0.050 | > 0.050 |
| Positive affect post-pre (mother) | 0.0007 | 0.004 | 14.68 | 0.19 | > 0.050 | > 0.050 |
| **Rules-based cooperative game** |  |  |  |  |  |  |
| Agreeableness (mother) | 0.0028 | 0.003 | 14.12 | 0.90 | > 0.050 | > 0.050 |
| Conscientiousness (mother) | 0.0039 | 0.002 | 13.97 | 1.68 | > 0.050 | > 0.050 |
| Extraversion (mother) | -0.0013 | 0.003 | 13.98 | -0.52 | > 0.050 | > 0.050 |
| Neuroticism (mother) | -0.0005 | 0.002 | 14.01 | -0.21 | > 0.050 | > 0.050 |
| Openness (mother) | 0.0004 | 0.002 | 14.05 | 0.19 | > 0.050 | > 0.050 |
| Effortful control (child) | 0.0003 | 0.003 | 13.97 | 0.11 | > 0.050 | > 0.050 |
| Negative affect (child) | -0.0035 | 0.003 | 14.01 | -1.39 | > 0.050 | > 0.050 |
| Surgency/extraversion (child) | -0.0002 | 0.005 | 13.98 | -0.05 | > 0.050 | > 0.050 |
| Negative affect post-pre (child) | -0.0005 | 0.006 | 14.11 | -0.09 | > 0.050 | > 0.050 |
| Positive affect post-pre (child) | -0.0018 | 0.004 | 14.23 | -0.46 | > 0.050 | > 0.050 |
| Negative affect post-pre (mother) | -0.0004 | 0.004 | 14.04 | -0.10 | > 0.050 | > 0.050 |
| Positive affect post-pre (mother) | 0.0001 | 0.005 | 14.11 | 0.02 | > 0.050 | > 0.050 |
| **Free interaction** |  |  |  |  |  |  |
| Agreeableness (mother) | 0.0034 | 0.001 | 14.41 | 2.27 | 0.039 | > 0.050 |
| Conscientiousness (mother) | 0.0001 | 0.001 | 13.38 | 0.06 | > 0.050 | > 0.050 |
| Extraversion (mother) | 0.0031 | 0.001 | 13.44 | 2.62 | 0.021 | > 0.050 |
| Neuroticism (mother) | 0.0036 | 0.001 | 13.45 | 3.20 | 0.007 | **0.021 \*** |
| Openness (mother) | 0.0003 | 0.001 | 13.87 | 0.27 | > 0.050 | > 0.050 |
| Effortful control (child) | -0.0018 | 0.001 | 13.42 | -1.32 | > 0.050 | > 0.050 |
| Negative affect (child) | 0.0003 | 0.001 | 13.46 | -0.28 | > 0.050 | > 0.050 |
| Surgency/extraversion (child) | 0.0060 | 0.002 | 13.35 | 2.50 | 0.026 | > 0.050 |
| Negative affect post-pre (child) | 0.0037 | 0.003 | 13.82 | 1.39 | > 0.050 | > 0.050 |
| Positive affect post-pre (child) | -0.0015 | 0.002 | 14.39 | -0.84 | > 0.050 | > 0.050 |
| Negative affect post-pre (mother) | -0.0027 | 0.002 | 13.37 | -1.39 | > 0.050 | > 0.050 |
| Positive affect post-pre (mother) | 0.0044 | 0.002 | 13.97 | 1.79 | > 0.050 | > 0.050 |

## 4. Discussion

The present study investigated mother–child INS using fNIRS hyperscanning by recording fronto-temporal brain activity from 33 dyads across three levels of interactivity: passive video co-exposure, a structured cooperative game, and unstructured free interaction. Prior to the experiment, mothers completed questionnaires assessing their own personality traits and their child’s temperament. In addition, mothers completed the PANAS before and after the experiment to assess post-pre changes in affect for both themselves and their child.

In the first part of the study, we assessed which ROI combinations were sensitive to co-presence and showed the hyperscanning effect (i.e., a statistically significant difference between true *vs*. surrogate dyads). We observed that social interactions with different levels of social interactivity elicited a hyperscanning effect mostly in ROI combinations involving the mother’s left IFG and the child’s right IFG. Notably, in our previous study (Carollo et al., 2025b), we found that during social interactions synchrony in the left IFG peaked during a cooperative Jenga task, whereas synchrony in the right IFG was maximal during video co-exposure, suggesting a task-dependent lateralization of INS. These findings are consistent with previous hyperscanning studies that have reported the involvement of inferior frontal regions during social interaction and interpersonal coordination (Liu et al., 2015; Nguyen et al., 2020; Reindl et al., 2018). Hyperscanning research has shown that parent–child cooperation elicits synchronized activity in prefrontal regions, including the dorsolateral prefrontal cortex and frontopolar cortex, and that this synchrony predicts cooperative performance (Reindl et al., 2018). Importantly, parent–child INS in these regions has been found to mediate the association between parental and child emotion regulation, suggesting that neural coupling may represent a key mechanism underlying socio-emotional development within the dyad (Reindl et al., 2018). At the single-brain level, the IFG is a key region in the understanding and imitation of others’ actions and intentions (Aziz-Zadeh et al., 2006; Carr et al., 2003; Iacoboni, 2005; Iacoboni and Dapretto, 2006; Lyons et al., 2006). In the parental brain, the IFG supports the perceptual-motor coupling and embodied simulation of the child’s action (Feldman, 2015). Notably, the IFG is also among the regions that exhibit selective and heightened responses to emotional infant faces in mothers compared to nulliparous women (Zhang et al., 2020). Furthermore, activity within the maternal left IFG has been shown to track the salience and rewarding value of child-related cues, with stronger responses when stimuli are perceived as one’s own child (Rigo et al., 2019). Hence, the hyperscanning effects observed in left maternal IFG–right child IFG pairs may reflect not only action understanding and mirroring processes, but also the selective motivational and affective significance of own-child cues during live interaction.

In the second part of the study analysis, we examined how dispositional and situational traits were related to INS levels. Only maternal neuroticism emerged as a significant predictor, with higher levels associated with increased INS. Neuroticism reflects a tendency to experience negative emotions, including anxiety and sadness (Vondra et al., 2006). Parents with higher levels of neuroticism and anxiety tend to be more likely to perceive their child’s environment as threatening and, as a consequence, exhibit parenting behaviors marked by lower sensitivity, over-protectiveness, and increased control (Bailes and Leerkes, 2021; Coplan et al., 2009). Similarly, in the current experiment, mothers with high neuroticism may remain highly vigilant to the child’s behaviors, resulting in tightly coordinated interactions in which mother and child continuously attend to and anticipate each other’s behavior. Consistent with this pattern, anxious caregivers have been shown to exhibit elevated physiological synchrony with their child (Smith et al., 2022) as well as altered, modality-specific behavioral coordination (Beebe et al., 2011), a profile that may reflect heightened vigilance rather than optimal co-regulation. At the neurobiological level, previous research suggests that increased mother-child synchrony may emerge in contexts of relational stress and maternal attachment anxiety, particularly during moments that require repair of coordination (Feldman et al., 2011). This heightened interpersonal coordination may reflect an overly stimulating interaction style that limits infants’ opportunities for self-regulation (Feldman, 2007; Smith et al., 2022). This pattern indicates that elevated interpersonal synchrony does not necessarily reflect optimal interpersonal exchanges (Gordon et al., 2024), but may instead signal heightened affiliative vigilance and increased sensitivity to the child’s needs and behaviors. In line with this, Nguyen et al. (2024) observed that maternal insecure attachment is associated with higher mother-child INS in a cooperative problem-solving task, suggesting that mothers with insecure attachment may engage in heightened efforts to coordinate and anticipate their child’s behavior.

Finally, when examining the effects of dispositional and situational traits and INS across the three levels of social interaction, neuroticism emerged as the only significant predictor in the video co-exposure and free interaction conditions, but not in the rules-based cooperative condition. These findings suggest that the association between neuroticism and INS might be contingent upon the degree of social interactivity characterizing the social task. In the video co-exposure and free interaction conditions, which are characterized by minimal behavioral constraints, mothers high in neuroticism may exhibit heightened vigilance and anticipatory monitoring of their child’s behavior. In these contexts, neuroticism might amplify INS and interpersonal synchrony, as anxious and closely attuned dyads monitor and respond continuously to each other’s signals (Beebe et al., 2011; Granat et al., 2017; Smith et al., 2022). In contrast, the absence of a significant association between maternal neuroticism and INS in the rule-based cooperative condition may be due to the structured nature of the task, which constrains behavior and reduces variability in interaction patterns (Krijnen et al., 2023), hence reducing the need for parental control and vigilance.

Notably, other predictors (i.e., including maternal personality traits beyond neuroticism, child temperament, and pre- and post-interaction affect) did not significantly explain variability in INS across conditions. This pattern suggests that relatively stable individual characteristics may play a more limited role in shaping neural synchrony during interaction than context-dependent regulatory processes. In particular, INS may be more strongly influenced by how the interaction unfolds in real time rather than by dispositional traits *per se*. In this sense, the quality and structure of the interactive exchange, the degree of mutual engagement, and the emotional availability within the dyad could overcome the effect of the other dispositional traits. From this perspective, neuroticism may have emerged as a significant predictor specifically because it is closely linked to heightened interpersonal vigilance and sensitivity to social cues, which become especially relevant in less structured, more open-ended interactional contexts.

### 4.1. Limitations and future studies

The results of this study should be interpreted in light of some limitations. Particularly, while the present study examined certain dispositional and situational characteristics, including the mother’s personality and the child’s temperament, it did not capture the full range of factors that may associates INS. For instance, factors such as maternal stress and the quality of mother-child attachment were not assessed in the study. In addition, the assessment of the traits considered here relied exclusively on self-report instruments, which may introduce subjective biases and limit the objectivity of the findings. Future studies could integrate multi-method approaches, such as behavioral observations and/or physiological measures, to complement the current findings.

In addition, the generalizability of the findings may be limited by the characteristics of the sample. Participants were drawn from a single cultural context and represented a relatively homogeneous group, which may restrict the extent to which the results can be generalized to more diverse populations. Moreover, the sample size may have been insufficient to reliably detect small effects, suggesting that future studies with larger and more heterogeneous samples are needed to strengthen the robustness and external validity of the findings. Regarding the homogeneity of the sample, the present study also focused on a relatively restricted developmental window, which may limit the understanding of how INS evolves across different stages of development. Future research could extend the age range of participating children in order to examine potential developmental changes in INS and to better capture how synchrony patterns emerge and transform over time.

A further limitation concerns the neural regions investigated. In this study, recordings were restricted to the bilateral IFG and TPJ, due to their involvement in INS. However, other brain areas, such as the medial prefrontal cortex, are also critically involved in social cognition and the coordination of interpersonal behavior and were not measured. This constraint was primarily due to the limited number of fNIRS optodes available. Expanding the coverage of neural recordings in future research would allow a more complete characterization of the neural networks underlying interpersonal synchrony and their interactions with individual and contextual factors.

Finally, the current study focused exclusively on mother–child dyads. Although mothers often represent the primary caregivers in early developmental contexts, fathers also play a crucial role in children’s socio-emotional development and interactive processes. Future studies should therefore consider including father–child dyads to explore whether similar or distinct patterns of INS emerge in paternal interactions.

## 5. Conclusion

In the current study, we conducted a secondary analysis of data from a previously published large-scale fNIRS hyperscanning project examining how interpersonal closeness and social interactivity relate to INS (Carollo et al., 2025b). Specifically, we investigated the contribution of dispositional and situational factors within 33 mother–child dyads across different interactive contexts. In this work, a significant hyperscanning effect emerged primarily in ROI combinations involving the mother’s left IFG and the child’s right IFG, suggesting that these regions are particularly sensitive to co-presence within this relational context. Given the role of the IFG in action understanding, embodied simulation, and parental responsiveness to emotionally salient own-child cues, this pattern likely reflects mirroring processes as well as motivational and affective salience in live mother–child interactions. Among the dispositional and situational variables examined, maternal neuroticism was the only statistically significant predictor of INS. Specifically, higher levels of maternal neuroticism were associated with increased INS, a pattern consistent with accounts linking anxiety-related traits to heightened parental vigilance and intensified monitoring of the child’s behavior. This positive association emerged during passive video co-exposure and free interaction, but not during the structured cooperative game, indicating that the influence of maternal dispositional traits may vary as a function of interactional demands and contextual structure. In less constrained contexts, heightened vigilance may amplify interpersonal coordination, whereas structured tasks may reduce behavioral variability and limit the expression of dispositional differences. Collectively, these findings underscore that INS is not a unitary marker of relational quality, but a dynamic, context-sensitive index shaped by both dispositional traits and interactional structure (Gordon et al., 2024). Importantly, our results, as well as emerging evidence (Nguyen et al., 2024), show that elevated INS should not be interpreted as unequivocally adaptive. Rather, it may reflect increased affiliative vigilance and tightly coupled interactional dynamics, potentially driven by heightened sensitivity to perceived environmental or relational threat.

## Authors contribution

Conceptualization: AC, AB, DD, SH, GE; Methodology: AC; Formal Analysis: AC, AB; Investigation: AC, DS; Writing–original draft preparation: AC, DS; Writing–review and editing: AC, AB, DS, DD, IG, GE, SH; Supervision: GE, SH. All authors have read and agreed to the published version of the manuscript.

