## Supplementary Materials 1, 2, 3 for "Over-Synchrony: Higher Maternal Neuroticism Associates with Stronger Interpersonal Neural Synchrony with Child During Passive and Free Interactions"

*Hyperscanning effect: true versus surrogate dyads*

Table S1 reports the results when comparing interpersonal neural synchrony computed on the deoxygenated hemoglobin (HbR) component between true and surrogate, artificially-paired dyads.

| **Fixed Effect** | **Estimate** | ***SE*** | ***df*** | ***t*-value** | ***p*-value** | **Adjusted *p*-value** | **Cohen's *d* 95% CI** |
|---|---|---|---|---|---|---|---|
| **Mother left IFG - Child left IFG** | | | | | | | |
| True dyads | 0.0008 | 0.0005 | 2207 | 1.56 | > 0.050 | > 0.050 | / |
| **Mother left IFG - Child right IFG** | | | | | | | |
| True dyads | 0.0016 | 0.0005 | 2220 | 3.99 | 0.003 | > 0.050 | / |
| **Mother left IFG - Child left TPJ** | | | | | | | |
| True dyads | 0.0016 | 0.0005 | 2143 | 2.97 | 0.003 | > 0.050 | / |
| **Mother left IFG - Child right TPJ** | | | | | | | |
| True dyads | 0.0015 | 0.0005 | 2221 | 2.86 | 0.004 | > 0.050 | / |
| **Mother right IFG - Child left IFG** | | | | | | | |
| True dyads | 0.0018 | 0.0005 | 2019 | 3.33 | < 0.001 | **0.028 *** | 0.15 [0.06, 0.24] |
| **Mother right IFG - Child right IFG** | | | | | | | |
| True dyads | 0.0019 | 0.0005 | 2052 | 3.72 | < 0.001 | **0.007 **** | 0.16 [0.08, 0.25] |
| **Mother right IFG - Child left TPJ** | | | | | | | |
| True dyads | 0.0013 | 0.0005 | 1984 | 2.32 | 0.020 | > 0.050 | / |
| **Mother right IFG - Child right TPJ** | | | | | | | |
| True dyads | 0.0008 | 0.0005 | 2047 | 1.56 | > 0.050 | > 0.050 | / |
| **Mother left TPJ - Child left IFG** | | | | | | | |
| True dyads | 0.0002 | 0.0005 | 2186 | 0.30 | > 0.050 | > 0.050 | / |
| **Mother left TPJ - Child right IFG** | | | | | | | |
| True dyads | 0.0010 | 0.0005 | 2235 | 1.95 | > 0.050 | > 0.050 | / |
| **Mother left TPJ - Child left TPJ** | | | | | | | |
| True dyads | 0.0015 | 0.0005 | 2136 | 2.72 | 0.007 | > 0.050 | / |
| **Mother left TPJ - Child right TPJ** | | | | | | | |
| True dyads | 0.0013 | 0.0005 | 2238 | 2.43 | 0.015 | > 0.050 | / |
| **Mother right TPJ - Child left IFG** | | | | | | | |
| True dyads | 0.0007 | 0.0006 | 1924 | 1.17 | > 0.050 | > 0.050 | / |
| **Mother right TPJ - Child right IFG** | | | | | | | |
| True dyads | 0.0015 | 0.0006 | 1935 | 2.51 | 0.012 | > 0.050 | / |
| **Mother right TPJ - Child left TPJ** | | | | | | | |
| True dyads | 0.0028 | 0.0006 | 1853 | 4.64 | < 0.001 | < **0.001 ***** | 0.22 [0.12, 0.31] |
| **Mother right TPJ - Child right TPJ** | | | | | | | |
| True dyads | 0.0013 | 0.0006 | 1939 | 2.20 | 0.028 | > 0.050 | / |

Table 1: Linear mixed-effects model results comparing true and surrogate mother–child dyads across inter-brain region pairs. Estimates reflect differences in interpersonal neural synchrony between true and randomly paired surrogate dyads for each mother–child region of interest combination (IFG = inferior frontal gyrus; TPJ = temporoparietal junction). Positive estimates indicate greater synchrony in true dyads relative to surrogate pairs. Reported statistics include fixed-effect estimates, standard errors (SE), degrees of freedom (df), $t$-values, unadjusted $p$-values, multiple-comparison–adjusted $p$-values, and effect sizes where applicable. Significant effects after correction are highlighted in bold. (* Adjusted $p$-value $< 0.050$; ** Adjusted $p$-value $< 0.010$; *** Adjusted $p$-value $< 0.001$)

*Association between dispositional and situational traits and interpersonal neural synchrony*

Table S2 reports the results when assessing the relationship between dispositional and situational traits and interpersonal neural synchrony computed on the HbR.

| **Predictor** | $\beta$ | ***SE*** | ***df*** | ***t*-value** | ***p*-value** |
|---|---|---|---|---|---|
| Agreeableness (mother) | 0.0020 | 0.002 | 14.25 | 1.16 | $> 0.050$ |
| Conscientiousness (mother) | 0.0004 | 0.001 | 13.92 | 0.30 | $> 0.050$ |
| Extraversion (mother) | 0.0007 | 0.001 | 13.93 | 0.53 | $> 0.050$ |
| Neuroticism (mother) | 0.0026 | 0.001 | 13.96 | 2.00 | $> 0.050$ |
| Openness (mother) | 0.0008 | 0.001 | 14.06 | 0.70 | $> 0.050$ |
| Effortful control (child) | 0.0008 | 0.002 | 13.92 | 0.50 | $> 0.050$ |
| Negative affect (child) | -0.0006 | 0.001 | 13.96 | -0.45 | $> 0.050$ |
| Surgency/extraversion (child) | 0.0010 | 0.003 | 13.91 | 0.36 | $> 0.050$ |
| Negative affect post-pre (child) | 0.0019 | 0.003 | 14.11 | 0.60 | $> 0.050$ |
| Positive affect post-pre (child) | -0.0005 | 0.002 | 14.22 | -0.23 | $> 0.050$ |
| Negative affect post-pre (mother) | -0.0004 | 0.002 | 13.97 | -0.16 | $> 0.050$ |
| Positive affect post-pre (mother) | 0.0006 | 0.003 | 14.12 | 0.21 | $> 0.050$ |

Table 2: Linear mixed-effects model examining dispositional and situational predictors of interpersonal neural synchrony. Maternal personality traits, child temperament dimensions, and post-pre changes in affect for both mother and child were entered as predictors of global interpersonal neural synchrony computed across region of interest pairs that previously demonstrated a significant hyperscanning effect. Reported statistics include regression coefficients ($\beta$), standard errors (SE), degrees of freedom (df), $t$-values, and associated $p$-values.

*Association between dispositional and situational traits and interpersonal neural synchrony across social interactivity levels*

Table S3 reports the results when assessing the relationship between dispositional and situational traits and interpersonal neural synchrony computed on the HbR by experimental condition.

| **Predictor** | $\beta$ | ***SE*** | ***df*** | ***t*-value** | ***p*-value** | **Adjusted *p*-value** |
|---|---|---|---|---|---|---|
| **Video co-exposure** | | | | | | |
| Agreeableness (mother) | -0.00025 | 0.002 | 15.46 | -0.10 | > 0.050 | > 0.050 |
| Conscientiousness (mother) | -0.00047 | 0.002 | 13.71 | -0.25 | > 0.050 | > 0.050 |
| Extraversion (mother) | 0.00096 | 0.002 | 13.75 | 0.47 | > 0.050 | > 0.050 |
| Neuroticism (mother) | 0.00379 | 0.002 | 13.81 | 1.98 | > 0.050 | > 0.050 |
| Openness (mother) | 0.00227 | 0.002 | 14.40 | 1.42 | > 0.050 | > 0.050 |
| Effortful control (child) | 0.00211 | 0.002 | 13.79 | 0.90 | > 0.050 | > 0.050 |
| Negative affect (child) | 0.00059 | 0.002 | 13.85 | 0.30 | > 0.050 | > 0.050 |
| Surgency/extraversion (child) | -0.00145 | 0.004 | 13.67 | -0.35 | > 0.050 | > 0.050 |
| Negative affect post-pre (child) | -0.00226 | 0.005 | 14.35 | -0.49 | > 0.050 | > 0.050 |
| Positive affect post-pre (child) | -0.00216 | 0.003 | 14.14 | -0.69 | > 0.050 | > 0.050 |
| Negative affect post-pre (mother) | 0.00032 | 0.003 | 13.84 | 0.09 | > 0.050 | > 0.050 |
| Positive affect post-pre (mother) | -0.00111 | 0.004 | 14.35 | -0.26 | > 0.050 | > 0.050 |
| **Rules-based cooperative game** | | | | | | |
| Agreeableness (mother) | 0.00369 | 0.002 | 14.25 | 1.54 | > 0.050 | > 0.050 |
| Conscientiousness (mother) | 0.00175 | 0.002 | 14.00 | 0.99 | > 0.050 | > 0.050 |
| Extraversion (mother) | -0.00015 | 0.002 | 14.01 | -0.08 | > 0.050 | > 0.050 |
| Neuroticism (mother) | 0.00241 | 0.002 | 14.07 | 1.34 | > 0.050 | > 0.050 |
| Openness (mother) | 0.0009 | 0.001 | 14.14 | 0.61 | > 0.050 | > 0.050 |
| Effortful control (child) | -0.00020 | 0.002 | 13.99 | -0.09 | > 0.050 | > 0.050 |
| Negative affect (child) | -0.00398 | 0.002 | 14.07 | -2.05 | > 0.050 | > 0.050 |
| Surgency/extraversion (child) | 0.00489 | 0.004 | 14.01 | 1.26 | > 0.050 | > 0.050 |
| Negative affect post-pre (child) | 0.00441 | 0.004 | 14.24 | 1.03 | > 0.050 | > 0.050 |
| Positive affect post-pre (child) | -0.00067 | 0.003 | 14.44 | -0.23 | > 0.050 | > 0.050 |
| Negative affect post-pre (mother) | 0.00008 | 0.003 | 14.12 | 0.03 | > 0.050 | > 0.050 |
| Positive affect post-pre (mother) | 0.00258 | 0.004 | 14.23 | 0.65 | > 0.050 | > 0.050 |
| **Free interaction** | | | | | | |
| Agreeableness (mother) | 0.00202 | 0.002 | 14.07 | 1.11 | > 0.050 | > 0.050 |
| Conscientiousness (mother) | 0.00014 | 0.001 | 13.42 | 0.10 | > 0.050 | > 0.050 |
| Extraversion (mother) | 0.00139 | 0.001 | 13.46 | 0.96 | > 0.050 | > 0.050 |
| Neuroticism (mother) | 0.00188 | 0.001 | 13.46 | 1.38 | > 0.050 | > 0.050 |
| Openness (mother) | -0.00045 | 0.001 | 13.73 | -0.40 | > 0.050 | > 0.050 |
| Effortful control (child) | -0.00002 | 0.002 | 13.44 | -0.01 | > 0.050 | > 0.050 |
| Negative affect (child) | 0.00094 | 0.001 | 13.47 | 0.64 | > 0.050 | > 0.050 |
| Surgency/extraversion (child) | 0.00024 | 0.003 | 13.40 | 0.08 | > 0.050 | > 0.050 |
| Negative affect post-pre (child) | 0.00261 | 0.003 | 13.69 | 0.81 | > 0.050 | > 0.050 |
| Positive affect post-pre (child) | 0.00093 | 0.002 | 14.07 | 0.42 | > 0.050 | > 0.050 |
| Negative affect post-pre (mother) | -0.00150 | 0.002 | 13.41 | -0.62 | > 0.050 | > 0.050 |
| Positive affect post-pre (mother) | -0.00015 | 0.003 | 13.81 | -0.05 | > 0.050 | > 0.050 |

Table 3: Linear mixed-effects model results examining whether maternal personality traits, child temperament dimensions, and post-pre affective changes predict interpersonal neural synchrony across region of interest pairs previously identified as showing a hyperscanning effect, separately for each experimental condition. Reported statistics include regression coefficients ($\beta$), standard errors (SE), degrees of freedom (df), $t$-values, unadjusted $p$-values, and multiple-comparison–adjusted $p$-values.
